# Antibody evolution to SARS-CoV-2 after single-dose Ad26.COV2.S vaccine

**DOI:** 10.1101/2022.03.31.486548

**Authors:** Alice Cho, Frauke Muecksch, Zijun Wang, Tarek Ben Tanfous, Justin DaSilva, Raphael Raspe, Brianna Johnson, Eva Bednarski, Victor Ramos, Dennis Schaefer-Babajew, Irina Shimeliovich, Juan Dizon, Kai-Hui Yao, Fabian Schmidt, Katrina G. Millard, Martina Turroja, Mila Jankovic, Thiago Y. Oliveira, Anna Gazumyan, Christian Gaebler, Marina Caskey, Theodora Hatziioannou, Paul D. Bieniasz, Michel C. Nussenzweig

## Abstract

The single dose Ad.26.COV.2 (Janssen) vaccine elicits lower levels of neutralizing antibodies and shows more limited efficacy in protection against infection than either of the available mRNA vaccines. In addition, the Ad.26.COV.2 has been less effective in protection against severe disease during the Omicron surge. Here, we examined the memory B cell response to single dose Ad.26.COV.2 vaccination. Compared to mRNA vaccines, Ad.26.COV.2 recipients had significantly lower numbers of RBD-specific memory B cells 1.5 or 6 months after vaccination. Memory antibodies elicited by both vaccine types show comparable neutralizing potency against SARS-CoV-2 and Delta. However, the number of memory cells producing Omicron neutralizing antibodies was somewhat lower after Ad.26.COV.2 than mRNA vaccination. The data help explain why boosting Ad.26.COV.2 vaccine recipients with mRNA vaccines is effective, and why the Janssen vaccine appears to have been less protective against severe disease during the Omicron surge than the mRNA vaccine.

**One-Sentence Summary:** Ad.26.COV.2 vaccine results in lower quantity but comparable quality of protective memory B cells compared to mRNA vaccines.

## Introduction

Severe Acute Respiratory Syndrome Coronavirus (SARS-CoV-2) produced a world-wide pandemic, infecting over 470 million people and is responsible for over 6 million deaths. In the United States, the FDA authorized the use of three vaccines encoding prefusion-stabilized SARS-CoV-2 spike: two mRNA-based, BNT162b2 from Pfizer-BioNTech and mRNA-1273 from Moderna, and an adenovirus-based vaccine, Ad26.COV2.S from Janssen (*1*). While both mRNA-based vaccines were initially approved as two-dose primary vaccine regimens, the replication-incompetent adenovirus (Ad) 26 vector-based Ad26.COV2.S vaccine received FDA emergency authorization as a single-dose vaccine. All three vaccines have since proven effective with substantial protection against COVID-19 infection, hospitalization, and death (*2, 3*).

However, protection against COVID-19 infection appeared to wane over time with Ad26.COV2.S demonstrating the most prominent decrease from 75% to 60% protective efficacy 5 months after vaccination, compared to a decrease in vaccine efficacy from 95% to either 67% or 80% after BNT162b2 and mRNA-1273 vaccination, respectively, over a similar period of time (*4*). Loss of protection against infection was associated with lower overall levels of SARS-CoV-2 spike protein (S)–specific antibodies and plasma neutralizing activity after Ad26.COV2.S immunization compared to mRNA vaccines (*5, 6*).

In contrast to protection from infection, Wuhan-Hu-1-based mRNA vaccines maintain effectiveness against hospitalization and death even in the face of infection with SARS-CoV-2 antigenic variants (*4, 7*). Protection from severe disease by mRNA vaccines is attributed in part to a diverse collection of memory B cells that develop cross reactivity against viral variants over time (*8*). Far less is known about the evolution of the memory B cell response after Ad.26.COV2 vaccination. Here, we report on memory B cell evolution over a 6-month period in a cohort of SARS-CoV-2-naïve individuals after Ad.26.COV2 immunization.

## Results

We studied the immune response to a single dose of the Ad.26.COV2.S (Janssen) vaccine in a cohort of 18 volunteers with no prior history of SARS-CoV-2 infection, recruited between April 26, 2021 and August 16, 2021 for sequential blood donations at 1.5 (median 46 days, range 27-72 days) and 6 months after vaccination (median 179 days, range 136-200 days). Volunteers ranged in age from 23-56 years and were 56% female and 44% male (for details, see Methods and table S1).

### Plasma binding and Neutralization

Plasma antibody binding titers to SARS-CoV-2 RBD were measured by enzyme-linked immunosorbent assays (ELISA) (*9, 10*). There was a 1.3-fold decrease in geometric mean IgG-binding titers against RBD between 1.5 and 6 months (p=0.07, Fig. 1a), which appears more modest than the 4.3-fold decrease reported for mRNA vaccinees at similar time points (*9*). RBD-binding IgG titers at the 1.5-month time point were comparable to a single dose of the mRNA vaccine and to convalescents 1.3 months after symptom onset (*9, 11*) (Fig. 1b). After 6 months, Janssen vaccine titers were significantly lower than in individuals who received 2 doses of an mRNA vaccine (*9*) (p=0.003, Fig 1b) but higher than convalescent infected individuals at a similar time after infection (*12*) (p=0.003, Fig. 1b). IgM and IgA-binding titers also decreased over time (p=0.01 and p=0.11, respectively, fig. S1a and b). IgM responses were comparable to both convalescent individuals and mRNA vaccinees, while IgA responses were significantly lower at both, 1.5- and 6-month time points when compared to mRNA vaccinees and convalescent individuals (fig. S1c and d).

**Fig. 1:**
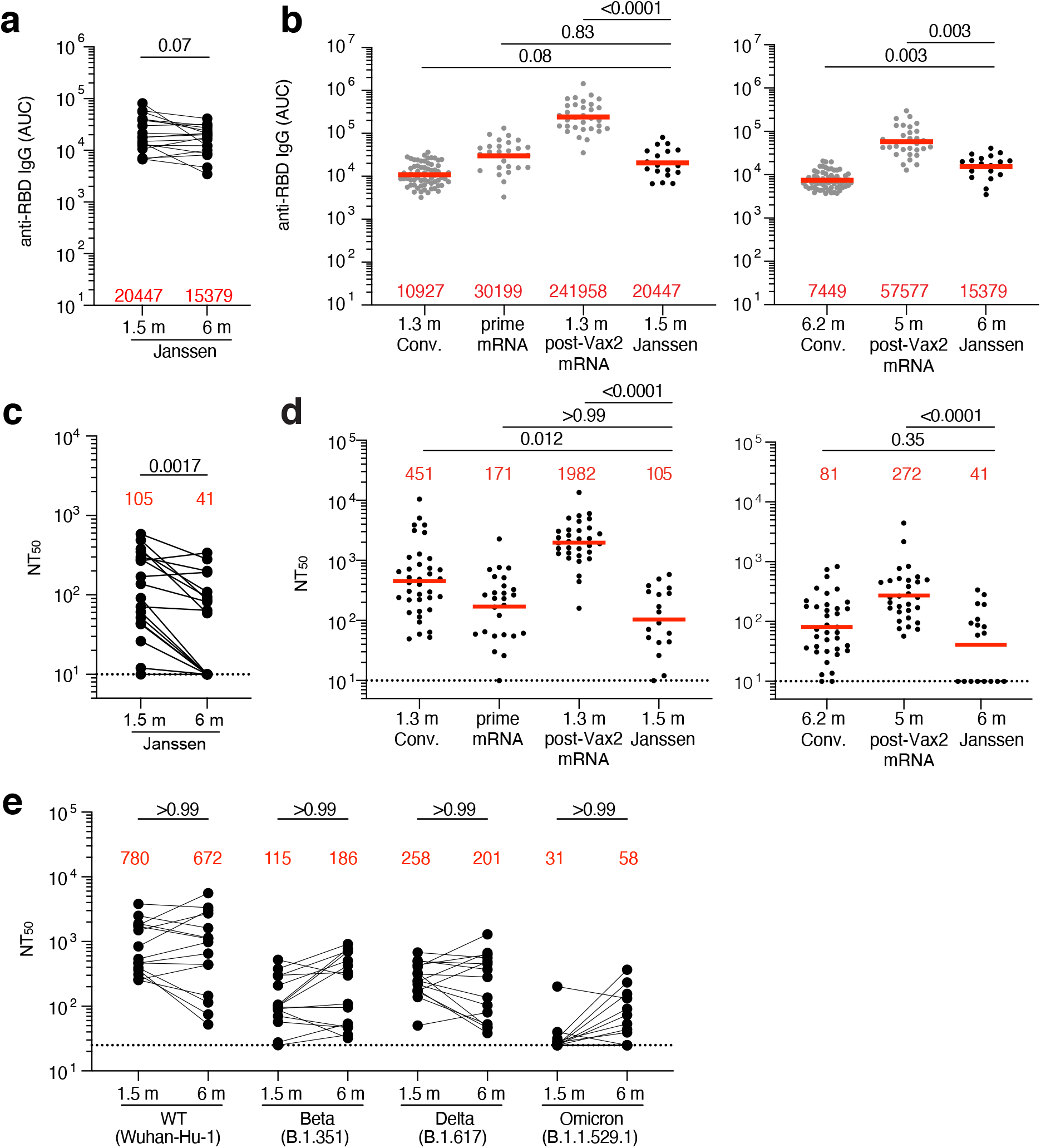
Plasma ELISAs and neutralizing activity. **a,** Graph shows area under the curve (AUC) for plasma IgG antibody binding to SARS-CoV-2 Wuhan-Hu-1 RBD 1.5 months (m) and 6 m post-vaccination for n=18 samples. Lines connect longitudinal samples. **b,** Graph shows AUC for plasma IgG binding to RBD in convalescent infected individuals 1.3 m post infection (*11*), and mRNA vaccinees after prime or 1.3 m post-second vaccination (Vax2) (*9*) compared to Janssen vaccinees 1.5 m post vaccination (left panel), or convalescent infected individuals 6.2 months post infection (*12*) and mRNA vaccinees 5 m post-Vax2 (*9*) compared to Janssen vaccinees at 6 m post vaccination (right panel). **c,** Graph shows anti-SARS-CoV-2 NT_50_s of plasma measured by a SARS-CoV-2 pseudotype virus neutralization assay using wild-type (Wuhan Hu-1 (*41*)) SARS-CoV-2 pseudovirus (*11, 42*) in plasma samples shown in panel **a. d,** NT_50_s of plasma measured by pseudotype virus neutralization assay comparing Janssen vaccinees to convalescent infected individuals (*11, 12*) and mRNA vaccinees (*9*) at either 1.5 months post-exposure (left panel) or 6 months post exposure (right), similar to plasma samples show in panel **b. e**, Plasma neutralizing activity against indicated SARS-CoV-2 variants of interest/concern for n=15 randomly selected samples assayed in HT1080Ace2 cl.14 cells. Wuhan-Hu-1 and Omicron BA.1 NT_50_ values are derived from (*43*). See Methods for a list of all substitutions/deletions/insertions in the spike variants. Deletions/substitutions corresponding to viral variants were incorporated into a spike protein that also includes the R683G substitution, which disrupts the furin cleavage site and increases particle infectivity. A corresponding WT control containing the R683G substitution was used in panel **e**. All experiments were performed at least in duplicate. Red bars and values represent geometric mean values. Statistical significance was determined by Wilcoxon matched-pairs signed rank test (**a** and **c**), two-tailed Kruskal-Wallis test with subsequent Dunn’s multiple comparisons (**b** and **d**) or Friedman test with subsequent Dunn’s multiple comparisons (**e**).

Neutralizing activity was determined for the same participants, using HIV-1 pseudotyped with Wuhan-Hu-1 SARS-CoV-2 Spike protein (*9, 10*) (table S1). 1.5 months after vaccination, individuals that received the Janssen vaccine had significantly lower neutralizing titers than either convalescents or vaccinees who received 2 doses of an mRNA vaccine (p=0.012 and p<0.0001, respectively, Fig. 1d). In contrast to reports that neutralizing titers increase marginally over time in Ad26.COV2.S vaccinees, there was a modest but significant 2.7-fold decrease in geometric mean neutralizing titers after 6 months in this cohort (*13–15*) (p=0.0017, Fig. 1c). As a result, 39% of the participants receiving the Janssen vaccine had neutralizing titers that were below the limit of detection in our assay (NT_50_<10). The neutralizing activity was comparable to convalescents but remained significantly lower than individuals who had received 2 doses of an mRNA vaccine (p<0.0001, Fig. 1d).

Plasma neutralizing activity for 15 randomly selected samples was also assessed against SARS-CoV-2 variants using pseudotype viruses with variant spikes (*9, 16*). Consistent with other reports (*13, 17*), at 1.5 months neutralizing titers against Beta, Delta, and Omicron were 6.8-, 3-, and 25-fold lower than Wuhan-Hu-1, respectively. In all cases the neutralizing titers were exceptionally low and did not change after 6 months (Fig. 1e).

### Memory B cell responses to SARS-CoV-2 RBD and NTD

Memory B cells contribute to long-term immune protection from serious disease by mediating rapid, anamnestic recall antibody responses (*18*). To examine the development of memory after Ad.26.COV2.S vaccination, we initially enumerated B cells expressing surface receptors binding to the Receptor-Binding Domain (RBD) or the N-Terminal Domain (NTD) of the SARS-CoV-2 spike protein using fluorescently labeled proteins (Fig. 2a, fig. S2a-c). The number of RBD-binding memory B cells at 1.5 months after Ad.26.COV2.S vaccination was significantly lower than for mRNA vaccinees 1.3 months after the second mRNA vaccine dose (p=0.008, Fig 2b, (*9*)). Although the number of RBD-binding memory cells increased between 1.5 and 6 months after the single-dose Janssen vaccine, the number remained lower than after mRNA vaccination at a similar time point (p=0.01, Fig. 2b). In contrast, the number of NTD binding memory B cells did not change between the 2 time points after Janssen vaccination and was significantly higher than after mRNA 5-6 months post vaccination (p=0.02, Fig. 2c). Additional phenotyping showed that RBD-specific memory B cells elicited by the Janssen vaccine showed the expected switch from IgM to IgG (fig. S2d).

**Fig. 2:**
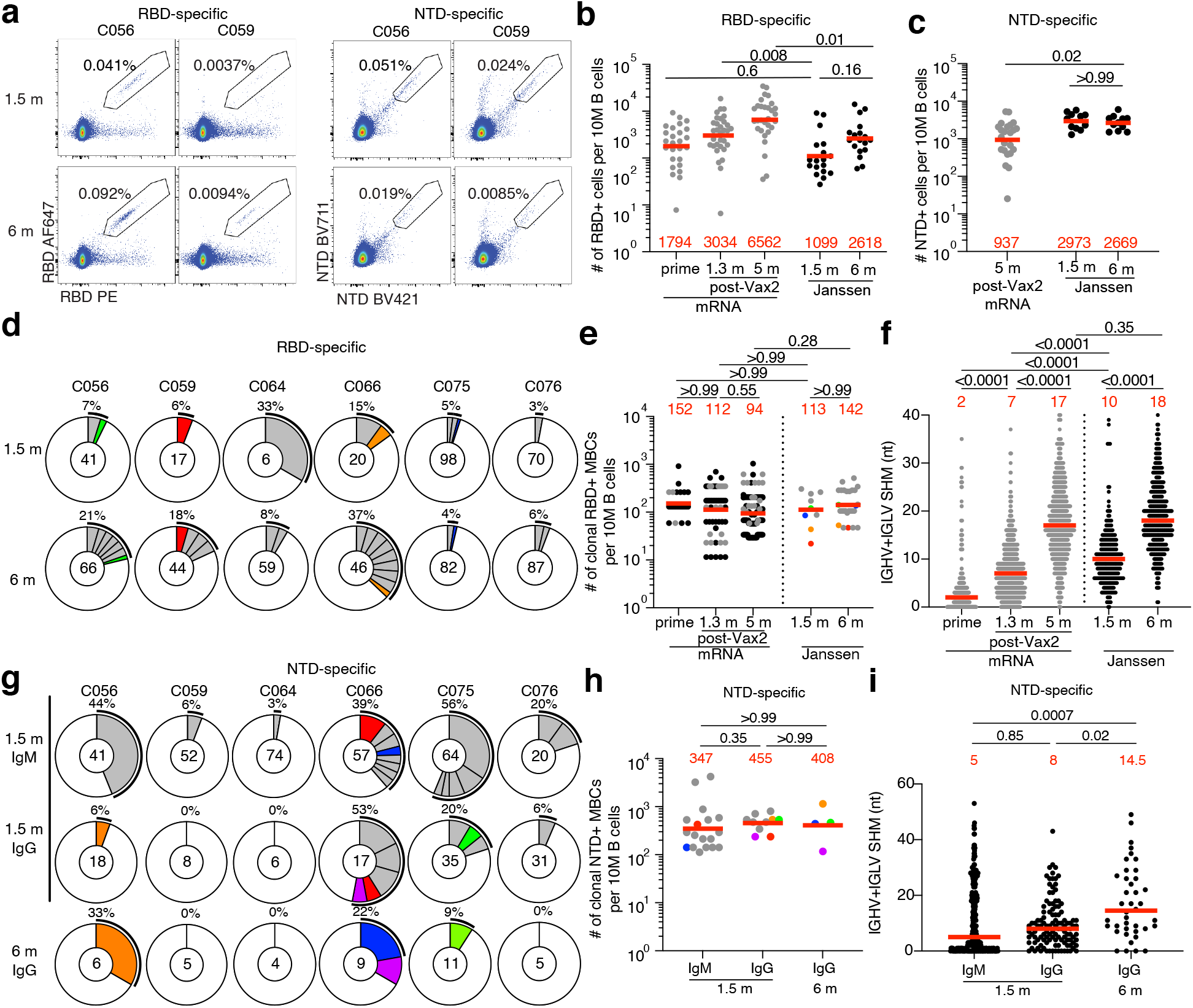
Anti-SARS-CoV-2 RBD and NTD B cells after vaccination. **a,** Representative flow cytometry plots showing dual AlexaFluor-647- and PE-Wuhan-Hu-1-RBD-binding (left panel) and BrilliantViolet-711- and BrilliantViolet-421-Wuhan-Hu-1 NTD-binding (right panel), single sorted B cells from 2 individuals at 1.5 months (m) or 6 m after vaccination. Gating strategy shown in fig. S2. Percentage of antigen-specific B cells is indicated. **b,** Graph summarizing the number of Wuhan-Hu-1 RBD-specific B cells per 10 million (M) B cells in Janssen vaccinees at 1.5 m and 6 m after vaccination (black dots, n=18) compared to mRNA vaccinees at prime, 1.3- and 5-m after Vax2 (*9*) (grey dots). **c,** Graph summarizing the number of Wuhan-Hu-1 NTD-specific B cells per 10 M B cells in Janssen vaccinees at 1.5 m and 6 m after vaccination (n=18), compared to mRNA vaccinees at 5 m after Vax2 (grey dots). **d,** Pie charts show the distribution of IgG antibody sequences obtained from Wuhan-Hu-1 RBD-specific memory B cells from 6 individuals after 1.5m and 6m post vaccination. Time points indicated to the left of the charts. The number inside the circle indicates the number of sequences analyzed for the individual denoted above the circle. Pie slice size is proportional to the number of clonally related sequences. The black outline and associated numbers indicate the percentage of clonally expanded sequences detected at each time point. Colored slices indicate persisting clones (same *IGHV* and *IGLV* genes with highly similar CDR3s, see Methods) found at more than one timepoint within the same individual. Grey slices indicate expanded clones unique to the timepoint. White slice represents sequences isolated only once. **e,** Graph shows the number of clonally expanded RBD-specific MBCs per 10 M B cell. Left panel represent clones from mRNA vaccinees after prime, or 1.3m and 5m post-Vax2 (black dots represent persisting clones, grey dots represent unique clones) (*8*). Right panel show clones from Janssen vaccinees at 1.5m or 6m post vaccination, with each dot representing one clone illustrated in Fig. 2d (color dots represent matched persisting clones, grey dots represent unique clones). **f,** Number of nucleotide somatic hypermutations (SHM) in *IGHV*+*IGLV* in RBD-specific sequences after 1.5- or 6-m post vaccination, compared to mRNA vaccinees (grey) after prime, or 1.3- and 5-m post-Vax2 (*9*). **g**, Pie charts showing distribution of IgM and IgG Wuhan-Hu-1 NTD-specific sequences after 1.5m and 6m post vaccination from same individuals as shown in Fig. 2d. Isotype and time point is indicated to left of graphs. **h,** Graph shows the number of clonally expanded NTD-specific MBCs per 10 M B cell, with each dot representing one clone illustrated in Fig. 2g (color dots represent matched persisting clones, grey dots represent unique clones). **i**, Number of nucleotide somatic hypermutations (SHM) in *IGHV* + *IGLV* in NTD-specific sequences after 1.5- or 6-m post-vaccination. Red bars and numbers in **b, c, e,** and **h,** represent geometric mean value (**b, c, e,** and **h)** or median values (**f** and **i)**. Statistical difference was determined by two-tailed Kruskal Wallis test with subsequent Dunn’s multiple comparisons (**b, c, e, f, h,** and **i**).

To examine the specificity and neutralizing activity of the antibodies produced by memory cells we purified single antigen-specific B cells, sequenced their antibody genes, and produced the recombinant antibodies *in vitro.* 636 paired anti-RBD antibody sequences were obtained from 6 vaccinees sampled at the 2 time points after Janssen vaccination (Fig. 2d, table S2). Clonally expanded RBD-specific B cells represented 6.3% and 13.5% of all memory cells at 1.5 and 6 months after vaccination, respectively, similar to mRNA vaccination (Fig. 2d and e). In addition, VH3-30 and VH3-53 genes were overrepresented to comparable degrees (fig. S3). However, there were very few clones that persisted over time (13% of all clonal expansions detected in Fig. 2d). The majority of expanded clones were found uniquely at one of the 2 time points (unique clones, 78%), suggesting ongoing memory B cell turnover (Fig. 2d). Continued memory B cell evolution was also evident in the accumulation of somatic mutations between 1.5 and 6 months (p<0.0001, Fig. 2f). Thus, although the absolute number of RBD-specific memory B cells 6 months after single dose Janssen was lower than after 2 doses of an mRNA vaccine, the two showed indistinguishable proportions of clonally expanded RBD-specific memory B cells that carry equivalent numbers of somatic mutations in their antibody genes.

To analyze the NTD-specific memory B cell repertoire, we sequenced 463 paired anti-NTD antibodies from the same 6 individuals (Fig. 2g, table S2). The geometric mean number of clonally expanded NTD specific memory cells was 4-fold greater than RBD-specific memory B cells after 1.5 months and remained 2.8-fold higher after 6 months (Fig. 2e and h). Similar to natural infection (*19*), VH4-39 and VH3-7 genes were over-represented in the NTD-specific memory B cell repertoire elicited by the Janssen vaccine (fig. S3). Expanded clones accounted for an average of 28% and 17% of the IgM and IgG repertoire 1.5 months after vaccination, respectively, and 13% of the IgG repertoire after 6 months. Like the RBD-specific memory B cells, only a minority (25%) of all expanded NTD-specific memory clones persisted between the two time points (Fig. 2g), and continued evolution was evident by accumulation of somatic mutations over time (p=0.02, Fig. 2i). In conclusion, the NTD-specific memory B cell compartment elicited by one dose of the Janssen vaccine is moderately larger in size and clonality to its anti-RBD counterpart.

### Neutralizing activity of monoclonal antibodies

192 anti-RBD monoclonal antibodies were expressed and tested for binding by ELISA. 93% (n=179) bound to the Wuhan-Hu-1 RBD, indicating the high efficiency RBD-specific memory B cell isolation (table S3). At the initial time point, the geometric mean ELISA half-maximal concentration (EC_50_) of the monoclonal antibodies obtained from Janssen vaccinees was significantly higher than individuals receiving a single dose of an mRNA vaccine (p=0.0001, Fig. 3a, (*9*)). However, the EC_50_ of RBD-binding antibodies elicited by the Janssen vaccine improved over time such that the antibodies elicited by the two vaccines had comparable EC_50_s after 5-6 months (Fig. 3a).

**Fig. 3:**
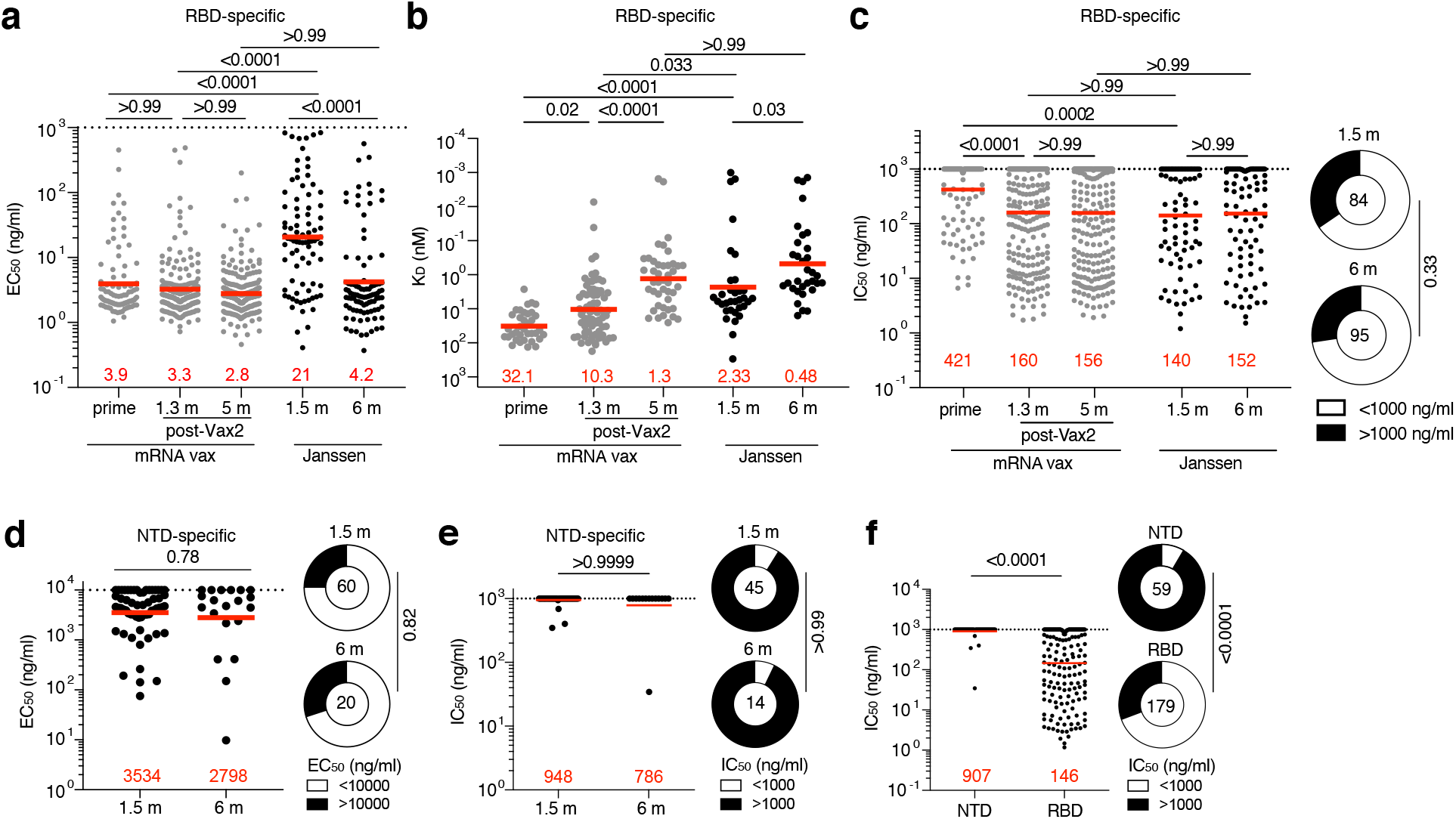
Anti-SARS-CoV-2 monoclonal antibodies. **a,** Graph shows half-maximal effective concentration (EC_50_) of n=179 Wuhan-Hu-1 RBD-binding monoclonal antibodies (mAbs) measured by ELISA against Wuhan-Hu-1 RBD 1.5m and 6m post vaccination, compared to EC_50_ measured in mRNA vaccinees after prime, 1.3- and 5-months post-Vax2 (*8, 9*). **b,** Graph showing affinity measurements (K_D_S) for Wuhan-Hu-1 RBD measured by BLI for antibodies cloned from mRNA vaccinees after prime, 1.3- and 6-months post-Vax2 (*8, 9*) compared to antibodies cloned from Janssen vaccinees at 1.5 m and 6 m (n=33, each) post vaccination. **c,** Graphs show anti-SARS-CoV-2 neutralizing activity of mAbs measured by a SARS-CoV-2 pseudotype virus neutralization assay using wild-type (Wuhan Hu-1 (*41*)) SARS-CoV-2 pseudovirus (*11, 42*) for antibodies cloned from mRNA vaccinees after prime, and 1.3- and 5-m post-Vax2 (*8, 9*) compared to antibodies cloned from Janssen vaccinees (n=179) at 1.5 m and 6 m post vaccination. Pie charts to the right indicated the frequency of neutralizing (IC_50_<1000 ng/mL, white) vs. non-neutralizing (IC_50_>1000 ng/mL, black) antibodies cloned from Janssen vaccinees. **d,** Graph showing EC_50_ of n=80 mAbs measured by ELISA against Wuhan-Hu-1 NTD after 1.5m and 6m post vaccination. Right panel shows pie charts indicating frequency of antibodies determined to bind (EC_50_<10000 ng/mL, white) or not bind (EC_50_>10000 ng/mL, black). **e**, Graph showing IC_50_ of NTD-specific antibodies at 1.5 m and 6 m post vaccination. Right panel shows pie charts indicating frequency of SARS-CoV-2 WT pseudovirus neutralizing (IC_50_<1000 ng/mL, white) vs. non-neutralizing (IC_50_>1000 ng/mL, black) NTD-specific mAbs. **f,** Graph comparing the IC_50_ of all NTD-specific mAbs (n=80) and RBD-specific mAbs (n=179) derived from Janssen vaccinees. Right panel shows pie charts indicating frequency of either NTD or RBD-specific neutralizing (IC_50_<1000 ng/mL, white) vs. non-neutralizing (IC_50_>1000 ng/mL, black) mAbs. Red bars and lines indicate geometric mean values. Statistical significance was determined by two-tailed Kruskal Wallis test with subsequent Dunn’s multiple comparisons (**a, b,** and **c)**, or by two-tailed Mann-Whitney test (**d, e,** and **f**). Pie charts were compared using a two-tailed Fisher’s exact test.

EC_50_s represent an indirect measure of affinity. To directly examine anti-RBD antibody affinity we performed biolayer interferometry (BLI) experiments on a subset of the antibodies (n=33 from 1.5 and 6 months, each). Affinity was significantly higher among antibodies elicited by the Janssen vaccine compared to those obtained after the mRNA prime and 2^nd^ dose (p<0.0001, and p=0.03, respectively, Fig. 3b, (*9*)). For both vaccine platforms, antibody affinity improved over time, reaching equivalent levels at the 5-6-month time point (Fig. 3b).

All 179 RBD-binding antibodies were tested for neutralization (84 and 95 antibodies isolated after 1.5 and 6 months, respectively). Compared to the mRNA prime, memory antibodies elicited by the Janssen vaccine were significantly more potent against viruses pseudotyped with the Wuhan-Hu-1 RBD (IC_50_ 140 vs 421 ng/ml, p=0.0002, Fig. 3c). However, the neutralizing activity of the anti-RBD memory antibodies elicited by mRNA vaccination improved after the second dose, and the two vaccines generated antibodies of equivalent potency after 5-6 months (IC_50_ 152 vs. 156, p>0.99, Fig 3c, (*9*)).

To examine the repertoire of NTD-specific memory B cells elicited by the Janssen vaccine, we expressed 60 and 20 antibodies obtained 1.5 and 6 months after vaccination, respectively (table S4). 59 bound to NTD with relatively poor EC_50_s that did not improve over time (Fig. 3d, table S4). When tested for neutralizing activity against Wuhan-Hu-1-pseudotyped virus, only 4 of the 59 NTD-binding monoclonal antibodies showed neutralizing activity, with no change over time (Fig. 3e). Thus, the overall frequency of memory B cells producing neutralizing anti-NTD antibodies is significantly lower than those producing anti-RBD (Fig. 3f). We conclude that anti-NTD memory antibodies are likely to make a more modest contribution to protection against subsequent viral challenge than their anti-RBD counterparts.

### Epitope specificity of RBD-binding antibodies

mRNA vaccination elicits anti-RBD antibodies that target 4 structurally defined classes of epitopes on the RBD (*8, 10, 20–22*). The relative distribution of epitopes targeted by RBD-binding antibodies can contribute to their potency and breadth. Whereas Class 1 and 2 antibodies, that block ACE2 binding directly, tend to be more potent, Class 3 and 4 target more conserved regions and can be broader (*8, 10, 12, 21*). To define the epitopes recognized by anti-RBD memory antibodies elicited by the Janssen vaccine, we performed BLI competition experiments. A preformed antibody-RBD-complex was exposed to a second antibody targeting one of four classes of structurally defined epitopes (*11, 20*) (C105 as Class 1; C144 as Class 2; C135 as Class 3; and C118 as Class 1/4). We examined 33 random RBD-binding antibodies obtained from the 1.5- and 6-month time points each, including 18 of 33 with IC_50_s lower than 1000 ng/mL. In contrast to the antibodies elicited after a single dose of an mRNA vaccine that primarily target Class 1 and 2 epitopes, Class 3 and 1/4 specific antibodies dominated the repertoire 1.5 months after Janssen vaccination (p= 0.016, Fig. 4a). This difference is particularly striking when considering neutralizing as opposed to non-neutralizing antibodies (Fig. 4b and c). However, shifts in the repertoire of the mRNA vaccinees over time alleviated these differences (Fig. 4a and b, (*8, 9*)).

**Fig. 4:**
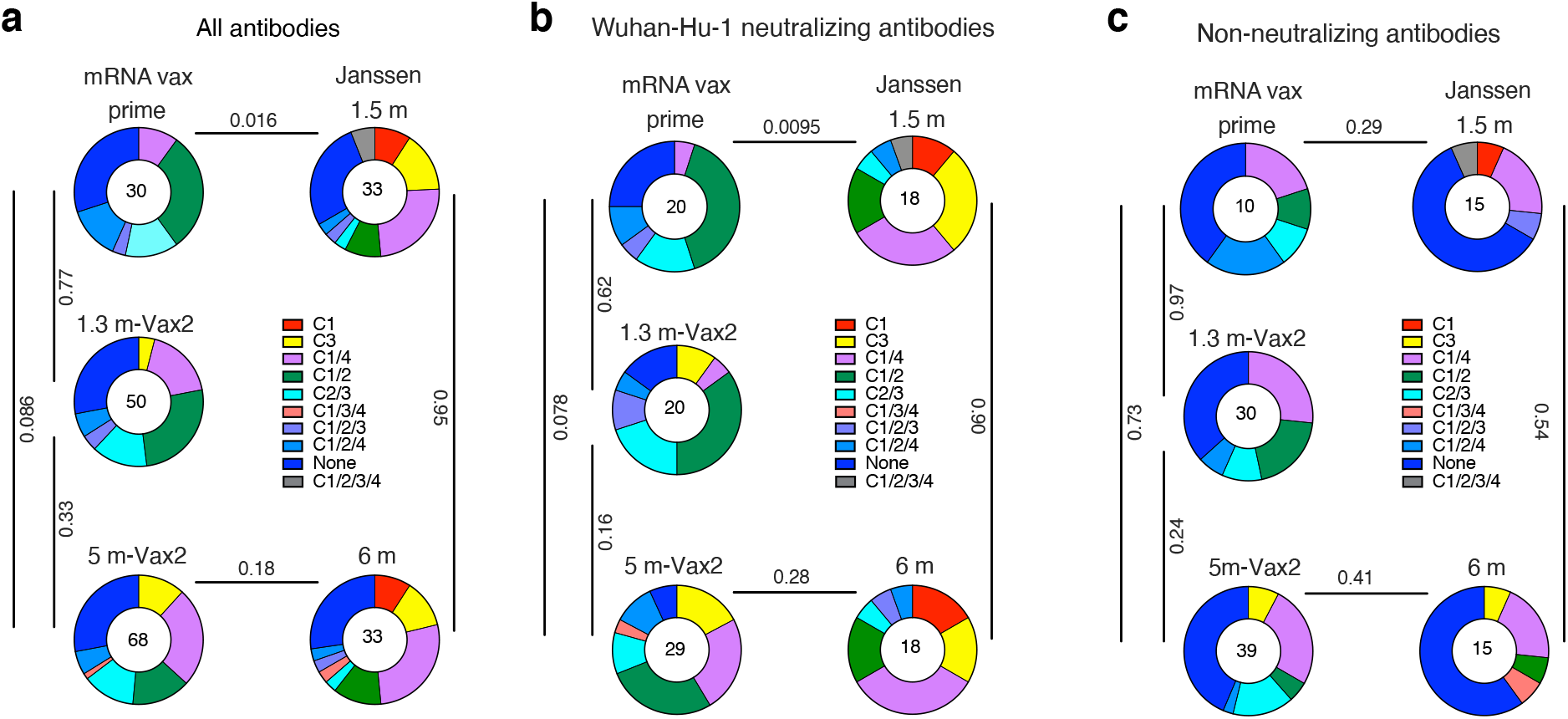
Epitope mapping. **a-c**, Results of epitope mapping performed by competition BLI, comparing mAbs cloned from Janssen vaccinees at 1.5 m and 6 m (n=33, each) post vaccination, to mAbs cloned from mRNA vaccinees at prime, or 1.3m and 5m post-Vax2 (*8, 9*). Pie charts show the distribution of the antibody classes among **a,** all RBD-binding antibodies, **b,** Wuhan-Hu-1 neutralizing antibodies only, or **c,** non-neutralizing antibodies only. Statistical significance was determined by using a two-tailed Chi-square test.

### Neutralizing Breadth

The neutralizing breadth of memory antibodies obtained from convalescent individuals increased significantly after 5 months (*10, 12, 21*). Memory antibodies elicited by mRNA vaccination show more modest improvement over the same period of time (*9*), which is further increased by a 3^rd^ dose (*8*). To determine how neutralizing breadth evolves after Janssen vaccination we analyzed a panel of 34 randomly selected Wuhan-Hu-1-neutralizing antibodies from Janssen vaccinees (n=16 at 1.5 months, and n=18 at 6 months). Neutralizing activity was measured against SARS-CoV-2 pseudoviruses carrying amino acid substitutions found in variants of concern. Neutralizing breadth improved significantly in Janssen vaccinees against pseudoviruses containing single amino acid substitutions found in different SARS-CoV-2 variants (K417N, N440K, and A475V, Fig. 5a, fig. S4a and b). These mutations typically alter the binding and neutralization properties of Class 1 and 3 antibodies (*21*).

**Fig. 5:**
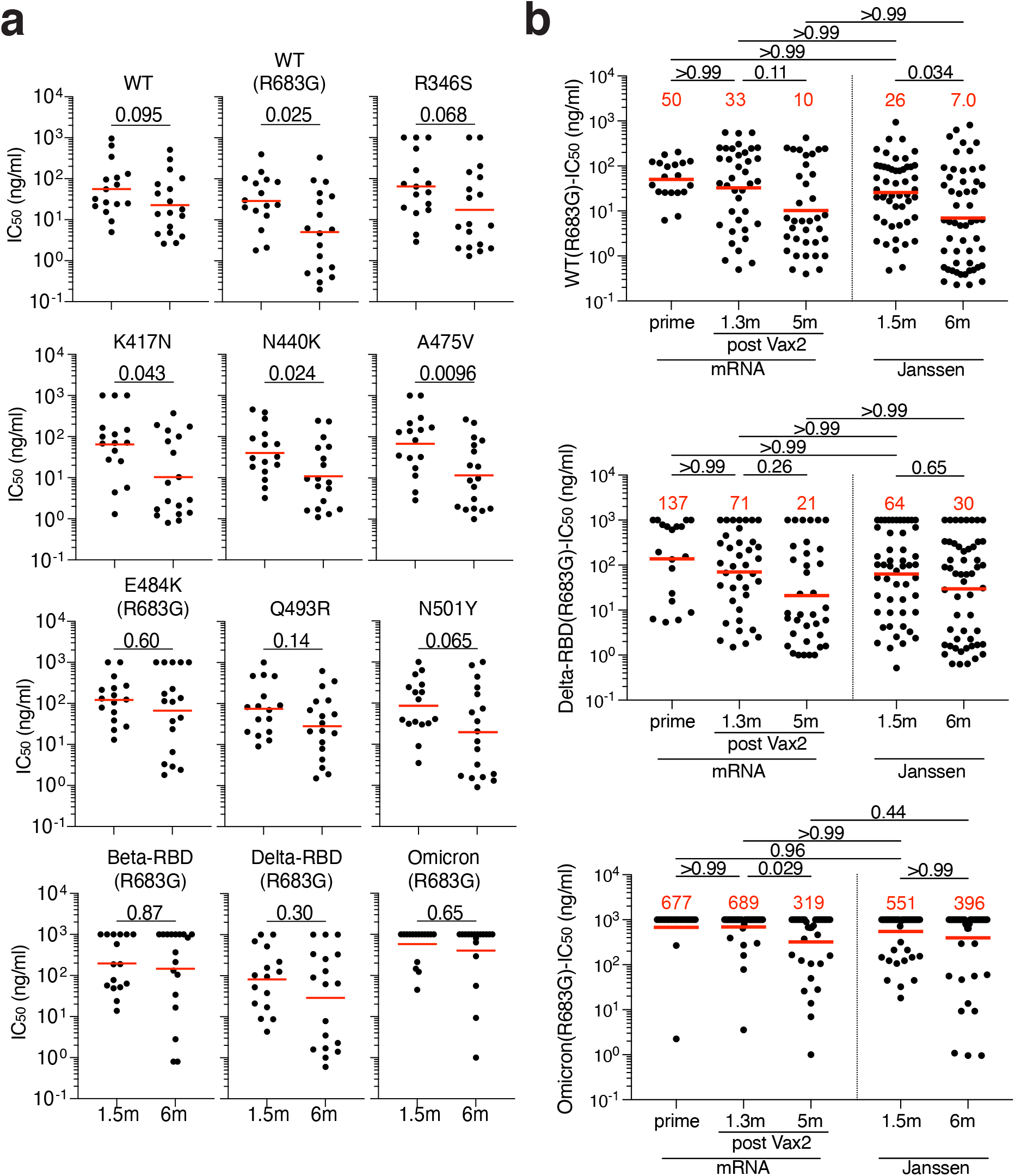
Breadth. **a,** Graphs showing IC_50_ neutralization activity of antibodies detected at 1.5 months (n=16) or 6 months (n=18) against indicated mutant SARS-CoV-2. **b,** Graphs showing IC_50_ neutralization activity of antibodies at 1.5 months (n=35) or 6 months (n=36) against wildtype (Wuhan-Hu-1 WT), Delta-RBD (L452R/T478K), and Omicron BA.1, compared to mRNA vaccinees at prime, and 1.3- and 5-m post-Vax2 (*8, 9*). The E484K, K417N/E484K/N501Y and L452R/T478K substitution, as well as the deletions/substitutions corresponding to viral variants, were incorporated into a spike protein that also includes the R683G substitution, which disrupts the furin cleavage site and increases particle infectivity. Neutralizing activity against mutant pseudoviruses were compared to a wildtype (WT) SARS-CoV-2 spike sequence (NC_045512), carrying R683G where appropriate. Red bars and lines indicated geometric mean values. Statistical significance in **a** was determined by two-tailed Mann-Whitney test, and in **b** by two-tailed Kruskal Wallis test with subsequent Dunn’s multiple comparisons.

A larger panel of randomly selected antibodies (n=71) with IC_50_s below 1000 ng/mL was tested for neutralizing activity against viruses pseudotyped with Wuhan-Hu-1, Delta, and Omicron RBDs (Fig. 5b and fig. S4c). In contrast to natural infection and mRNA vaccination there was no improvement in neutralizing activity against Delta or Omicron between 1.5 and 6 months after Janssen vaccination. Nevertheless, 86% of the 6-month memory antibodies tested neutralized Delta and 31% neutralized Omicron (fig. S4c). Thus, 6 months after vaccination the memory B cell compartment in Ad26.COV2.S recipients is smaller in size than the RBD-specific memory B cell compartment in mRNA vaccinees but contains cells with the ability to produce antibodies with comparable activity against Delta and Omicron.

## Discussion

Administration of a single dose of the Ad26.COV2.S vaccine results in less effective protection against infection than mRNA vaccination, and also affords lower levels of protection against severe disease and hospitalization from COVID-19 (*4, 6, 23*). The difference in protective efficacy from infection between the 2 vaccine modalities has been attributed to significantly lower levels of circulating neutralizing antibodies elicited by the Janssen vaccine (*14, 24*). We find that 5-6 months after vaccination there is a 2.5-fold difference in the number of memory B cells produced by the 2 vaccine modalities. A third mRNA dose further magnifies the difference to nearly 6 fold (*8*). Nevertheless, the antibodies encoded by the individual memory cells show similar levels of activity against Wuhan-Hu-1, and Delta, and Omicron BA.1. The ability of these cells to respond rapidly to viral challenge may account in part for the partial protection against severe disease by Ad26.COV2.S vaccination.

Circulating antibodies are produced from plasma cells that are selected in germinal centers and extrafollicular foci from a diverse cohort of follicular B cells based primarily on their affinity for antigen (*25, 26*). Many of the plasma cells produced during the early stages of the immune response are short-lived resulting in a transient early peak in circulating antibody levels (*27*). Memory B cells develop in the same two microanatomic compartments, but their development is regulated by an entirely different cellular and molecular program (*18, 28–31*). As a result, memory B cells are long-lived and express a diverse collection of antibodies with differing affinities, neutralizing activity, and breath (*31, 32*).

The relatively poor plasma binding and neutralizing titers elicited by the Ad26.COV2.S vaccine compared to mRNA vaccines points to more modest elicitation of plasma cell responses by Ad26.COV2.S. In addition, the number of memory B cells elicited by the single dose Ad26.COV2.S vaccine is smaller than 2 doses of the mRNA vaccines at all time points examined. A third mRNA booster vaccination amplifies this difference. However, neutralizing potency and breadth develop rapidly after Ad26.COV2.S vaccination and the memory antibodies elicited by the two vaccine modalities display comparable potency and breadth against Wuhan-Hu-1, and Delta at both, 1.5 and 6 months after vaccination. Activity against Omicron was lower after Ad26.COV2.S but the difference was not statistically significant.

Class 1 and 2 antibodies develop early after infection or mRNA immunization and are generally more potent than class 3 and 4, because they interfere directly with the interaction between the SARS-CoV-2 RBD and its cellular receptor ACE2 (*10, 20, 21*). However, this renders Class 1 and 2 antibodies highly sensitive to amino acid substitutions within the ACE2 binding ridge of the RBD found in many SARS-CoV-2 variants (*10, 21*). The epitopes targeted by Class 3 and 4 are generally more conserved and antibodies binding to these epitopes may be more broadly reactive. Class 3 and 4 antibodies develop earlier in Ad26.COV2.S than in mRNA vaccinees, leading to a more diverse early B cell memory response. Nevertheless, continued evolution is a feature of memory B cell responses to both vaccine modalities, and they become comparable in this respect after 5-6 months.

Neutralizing antibodies are the best correlate of protection, and when provided early they are also therapeutic against COVID-19 (*33–37*). Although memory B cells are quiescent and do not contribute to the pool of circulating antibodies under steady state conditions, they can be recalled rapidly upon challenge to develop into antibody producing cells (*38, 39*). Our observations show that a diverse memory B cell compartment develops in response to the Ad26.COV2.S vaccine including a subset of cells that express antibodies that potently neutralize antigenically divergent variants. Rapid activation of these cells and antibody production upon SARS-CoV-2 infection may explain why the Ad26.COV2.S vaccine is partially effective at providing protection against severe disease following breakthrough infection, and priming with this vaccine supports robust responses after heterologous boosting with mRNA vaccines (*14, 23, 40*).

## Acknowledgements

We thank all study participants who devoted time to our research, The Rockefeller University Hospital nursing staff and Clinical Research Support Office. We thank all members of the M.C.N. laboratory for helpful discussions, Maša Jankovic and Gabriel Scrivanti for laboratory support and Kristie Gordon for technical assistance with cell-sorting experiments.

## Author information

AC, FM, and ZW contributed equally to this work.

## Author Contributions

AC, FM, ZW, TH, PDB, and MCN. conceived, designed, and analyzed the experiments. MC and CG designed clinical protocols. AC, FM, ZW, TBT, JD, RR, EB, DS-B, KY, MJ, and FS carried out experiments. BJ and AG produced antibodies. MT, KGM, IS, JD, CG and MC recruited participants, executed clinical protocols, and processed samples. TYO and VR performed bioinformatic analysis. AC, FM, ZW, CG, TH, PDB, and MCN wrote the manuscript with input from all co-authors.

## Corresponding authors

Correspondence should be addressed to Theodora Hatziioannou, Paul D. Bieniasz, or Michel C. Nussenzweig.

## Data and materials availability

Data are provided in Tables S1-S4. The raw sequencing data and computer scripts associated with Fig. 2 have been deposited at Github (https://github.com/stratust/igpipeline/tree/igpipeline2_timepoint_v2). This study also uses data from “A Public Database of Memory and Naive B-Cell Receptor Sequences” (https://doi.org/10.5061/dryad.35ks2), PDB (6VYB and 6NB6), cAb-Rep (https://cab-rep.c2b2.columbia.edu/), Sequence Read Archive (accession SRP010970), and from “High frequency of shared clonotypes in human B cell receptor repertoires” (https://doi.org/10.1038/s41586-019-0934-8). Computer code to process the antibody sequences is available at GitHub (https://github.com/stratust/igpipeline/tree/igpipeline2_timepoint_v2).

## Materials and Methods

### Study participants

Participants were healthy volunteers who had previously received one dose of the Janssen (Ad26.COV2.S) vaccine against wildtype (Wuhan-Hu-1) strain of the severe acute respiratory syndrome coronavirus 2 (SARS-CoV-2). For this study, participants were recruited for serial blood donations at the Rockefeller University Hospital in New York between April 26, 2021 and August 16, 2021. Eligible participants (n=18) were healthy adults with no history of infection with SARS-CoV-2 during or prior to the observation period (as determined by clinical history and confirmed through serology testing) who had received only one dose the SARS-CoV-2 Janssen Ad26.COV2.S vaccine. Exclusion criteria include presence of clinical signs and symptoms suggestive of acute infection with or a positive reverse transcription polymerase chain reaction (RT-PCR) results for SARS-CoV-2 in saliva, or a positive COVID-19 (coronavirus disease 2019) serology. Participants presented to the Rockefeller University Hospital for blood sample collection and were asked to provide details of their vaccination regimen, possible side effects, comorbidities, and possible COVID-19 history. Clinical data collection and management were carried out using the software iRIS by iMedRIS (v. 11.02). All participants provided written informed consent before participation in the study and the study was conducted in accordance with Good Clinical Practice. The study was performed in compliance with all relevant ethical regulations and the protocol (DRO-1006) for studies with human participants was approved by the Institutional Review Board of the Rockefeller University. For detailed participant characteristics see table S1.

### Blood samples processing and storage

Peripheral Blood Mononuclear Cells (PBMCs) obtained from samples collected at Rockefeller University were purified as previously reported by gradient centrifugation and stored in liquid nitrogen in the presence of Fetal Calf Serum (FCS) and Dimethylsulfoxide (DMSO) (*11, 12*). Heparinized plasma and serum samples were aliquoted and stored at −20°C or less. Prior to experiments, aliquots of plasma samples were heat-inactivated (56°C for 30 minutes) and then stored at 4°C.

### ELISAs

Enzyme-Linked Immunosorbent Assays (ELISAs) (*44, 45*) to evaluate antibodies binding to SARS-CoV-2 RBD or NTD were performed by coating of high-binding 96-half-well plates (Corning 3690) with 50 μl per well of a 1μg/ml protein solution in Phosphate-buffered Saline (PBS) overnight at 4°C. Plates were washed 6 times with washing buffer (1× PBS with 0.05% Tween-20 (Sigma-Aldrich)) and incubated with 170 μl per well blocking buffer (1× PBS with 2% BSA and 0.05% Tween-20 (Sigma)) for 1 hour at room temperature. Immediately after blocking, monoclonal antibodies or plasma samples were added in PBS and incubated for 1 hour at room temperature. Plasma samples were assayed at a 1:66 starting dilution and 10 additional threefold serial dilutions. Monoclonal antibodies were tested at 10 μg/ml starting concentration and 10 additional fourfold serial dilutions. Plates were washed 6 times with washing buffer and then incubated with anti-human IgG, IgM or IgA secondary antibody conjugated to horseradish peroxidase (HRP) (Jackson ImmunoResearch 109-036-088, 109-035-129, and Sigma A0295) in blocking buffer at a 1:5,000 dilution (IgM and IgG) or 1:3,000 dilution (IgA). Plates were developed by addition of the HRP substrate, 3,3’,5,5’-Tetramethylbenzidine (TMB) (ThermoFisher) for 10 minutes (plasma samples) or 4 minutes (monoclonal antibodies). The developing reaction was stopped by adding 50 μl of 1 M H_2_SO_4_ and absorbance was measured at 450 nm with an ELISA microplate reader (FluoStar Omega, BMG Labtech) with Omega and Omega MARS software for analysis. For plasma samples, a positive control (plasma from participant COV72, diluted 66.6-fold and ten additional threefold serial dilutions in PBS) was added to every assay plate for normalization. The average of its signal was used for normalization of all the other values on the same plate with Excel software before calculating the area under the curve using Prism V9.1 (GraphPad). Negative controls of pre-pandemic plasma samples from healthy donors were used for validation (for more details, please see (*11*)). For monoclonal antibodies, the ELISA half-maximal concentration (EC_50_) was determined using four-parameter nonlinear regression (GraphPad Prism V9.1). EC_50_s above 1000 ng/mL for RBD-binding were considered non-binders; EC_50_s above 10000 ng/mL for NTD-binding were considered non-binders.

### Proteins

The mammalian expression vector encoding the Receptor Binding-Domain (RBD) of SARS-CoV-2 (GenBank MN985325.1; Spike (S) protein residues 319-539) was previously described (*46*). Mammalian expression vector encoding the SARS-CoV-2 Wuhan-Hu-1 NTD (GenBank MN985325.1; S protein residues 14-307) was previously described (*19*).

### SARS-CoV-2 pseudotyped reporter virus

A panel of plasmids expressing RBD-mutant SARS-CoV-2 spike proteins in the context of pSARS-CoV-2-S_Δ19_ has been described (*9, 16, 21, 47*). Variant pseudoviruses resembling SARS-CoV-2 variants Beta (B.1.351), B.1.526, Delta (B.1.617.2) and Omicron BA.1 (B.1.1.529) have been described before (*9, 10, 43*) and were generated by introduction of substitutions using synthetic gene fragments (IDT) or overlap extension PCR mediated mutagenesis and Gibson assembly. Specifically, the variant-specific deletions and substitutions introduced were:

Beta: D80A, D215G, L242H, R246I, K417N, E484K, N501Y, D614G, A701V
Delta: T19R, Δ156-158, L452R, T478K, D614G, P681R, D950N
Omicron BA.1: A67V, Δ69-70, T95I, G142D, Δ143-145, Δ211, L212I, ins214EPE, G339D, S371L, S373P, S375F, K417N, N440K, G446S, S477N, T478K, E484A, Q493K, G496S, Q498R, N501Y, Y505H, T547K, D614G, H655Y, H679K, P681H, N764K, D796Y, N856K, Q954H, N969H, N969K, L981F

The E484K, K417N/E484K/N501Y and L452R/T478K substitution, as well as the deletions/substitutions corresponding to variants of concern listed above, were incorporated into a spike protein that also includes the R683G substitution, which disrupts the furin cleavage site and increases particle infectivity. Neutralizing activity against mutant pseudoviruses were compared to a wildtype (WT) SARS-CoV-2 spike sequence (NC_045512), carrying R683G where appropriate.

SARS-CoV-2 pseudotyped particles were generated as previously described (*11, 42*). Briefly, 293T (CRL-11268) cells were obtained from ATCC, and the cells were transfected with pNL4-3 DEnv-nanoluc and pSARS-CoV-2-S_Δ19_. Particles were harvested 48 hours post-transfection, filtered and stored at −80°C.

### Pseudotyped virus neutralization assay

Four-to five-fold serially diluted pre-pandemic negative control plasma from healthy donors, plasma from individuals who received Ad26.COV2.S vaccines, or monoclonal antibodies were incubated with SARS-CoV-2 pseudotyped virus for 1 hour at 37 °C. The mixture was subsequently incubated with 293T_Ace2_ cells (*11*) (for all WT neutralization assays) or HT1080Ace2 cl14 (for all mutant panels and variant neutralization assays) cells (*16*) for 48 hours after which cells were washed with PBS and lysed with Luciferase Cell Culture Lysis 5× reagent (Promega). Nanoluc Luciferase activity in lysates was measured using the Nano-Glo Luciferase Assay System (Promega) with the Glomax Navigator (Promega) or ClarioStar multi-mode microplate reader (BMG). The relative luminescence units were normalized to those derived from cells infected with SARS-CoV-2 pseudotyped virus in the absence of plasma or monoclonal antibodies. The half-maximal neutralization titers for plasma (NT50) or half-maximal and 90% inhibitory concentrations for monoclonal antibodies (IC_50_ and IC90) were determined using four-parameter nonlinear regression (least squares regression method without weighting; constraints: top=1, bottom=0) (GraphPad Prism).

### Biotinylation of viral protein for use in flow cytometry

Purified and Avi-tagged SARS-CoV-2 Wuhan-Hu-1 RBD and NTD were biotinylated using the Biotin-Protein Ligase-BIRA kit according to manufacturer’s instructions (Avidity) as described before (*11*). Ovalbumin (Sigma, A5503-1G) was biotinylated using the EZ-Link Sulfo-NHS-LC-Biotinylation kit according to the manufacturer’s instructions (Thermo Scientific). Biotinylated ovalbumin was conjugated to streptavidin-BB515 (BD, 564453). RBD was conjugated to streptavidin-PE (BD Biosciences, 554061) and streptavidin-AF647 (Biolegend, 405237) (*11*). NTD was conjugated to streptavidin-BV421 (Biolegend, 405225) and streptavidin-BV711 (BD Biosciences, 563262).

### Flow cytometry and single cell sorting

Single-cell sorting by flow cytometry was described previously (*11*). Briefly, peripheral blood mononuclear cells were enriched for B cells by negative selection using a pan-B-cell isolation kit according to the manufacturer’s instructions (Miltenyi Biotec, 130-101-638). The enriched B cells were incubated in Flourescence-Activated Cell-sorting (FACS) buffer (1× PBS, 2% FCS, 1 mM ethylenediaminetetraacetic acid (EDTA)) with the following anti-human antibodies (all at 1:200 dilution): anti-CD20-PECy7 (BD Biosciences, 335793), anti-CD3-APC-eFluro780 (Invitrogen, 47-0037-41), anti-CD8-APC-eFluor780 (Invitrogen, 47-0086-42), anti-CD16-APC-eFluor780 (Invitrogen, 47-0168-41), anti-CD14-APC-eFluor780 (Invitrogen, 47-0149-42), as well as Zombie NIR (BioLegend, 423105) and fluorophore-labeled Wuhan-Hu-1 RBD, NTD, and ovalbumin (Ova) for 30 min on ice. AccuCheck Counting Beads (Life Technologies, PCB100) were added to each sample according to manufacturer’s instructions. Single CD3-CD8-CD14-CD16-CD20+Ova-B cells that were either RBD-PE+RBD-AF647+ or NTD-BV711+NTD-BV421+ were sorted into individual wells of 96-well plates containing 4 μl of lysis buffer (0.5× PBS, 10 mM Dithiothreitol (DTT), 3,000 units/ml RNasin Ribonuclease Inhibitors (Promega, N2615)) per well using a FACS Aria III and FACSDiva software (Becton Dickinson) for acquisition and FlowJo for analysis. The sorted cells were frozen on dry ice, and then stored at −80 °C or immediately used for subsequent RNA reverse transcription. For B cell phenotype analysis, in addition to above antibodies, B cells were also stained with following anti-human antibodies (all at 1:200 dilution): anti-IgD-BV650 (BD, 740594), anti-CD27-BV786 (BD biosciences, 563327), anti-CD19-BV605 (Biolegend, 302244), anti-CD71-PerCP-Cy5.5 (Biolegend, 334114), anti-IgG-PECF594 (BD, 562538), anti-IgM-AF700 (Biolegend, 314538), anti-IgA-Viogreen (Miltenyi Biotec, 130-113-481).

### Antibody sequencing, cloning and expression

Antibodies were identified and sequenced as described previously (*11, 48*). In brief, RNA from single cells was reverse-transcribed (SuperScript III Reverse Transcriptase, Invitrogen, 18080-044) and the cDNA was stored at −20 °C or used for subsequent amplification of the variable IGH, IGL and IGK genes by nested PCR and Sanger sequencing. Sequence analysis was performed using MacVector. Amplicons from the first PCR reaction were used as templates for sequence- and ligation-independent cloning into antibody expression vectors. Recombinant monoclonal antibodies were produced and purified as previously described (*11*).

### Biolayer interferometry

Biolayer interferometry assays were performed as previously described (*11*). Briefly, we used the Octet Red instrument (ForteBio) at 30°C with shaking at 1,000 r.p.m. Epitope binding assays were performed with protein A biosensor (ForteBio 18-5010), following the manufacturer’s protocol “classical sandwich assay” as follows: (1) Sensor check: sensors immersed 30 sec in buffer alone (buffer ForteBio 18-1105), (2) Capture 1st Ab: sensors immersed 10 min with Ab1 at 10 μg/mL, (3) Baseline: sensors immersed 30 sec in buffer alone, (4) Blocking: sensors immersed 5 min with IgG isotype control at 10 μg/mL. (5) Baseline: sensors immersed 30 sec in buffer alone, (6) Antigen association: sensors immersed 5 min with RBD at 10 μg/mL. (7) Baseline: sensors immersed 30 sec in buffer alone. (8) Association Ab2: sensors immersed 5 min with Ab2 at 10 μg/mL. Curve fitting was performed using the Fortebio Octet Data analysis software (ForteBio).

### Computational analyses of antibody sequences

Antibody sequences were trimmed based on quality and annotated using Igblastn v.1.14. with IMGT domain delineation system. Annotation was performed systematically using Change-O toolkit v.0.4.540 (*49*). Clonality of heavy and light chain was determined using DefineClones.py implemented by Change-O v0.4.5 (*49*). The script calculates the Hamming distance between each sequence in the data set and its nearest neighbor. Distances are subsequently normalized and to account for differences in junction sequence length, and clonality is determined based on a cut-off threshold of 0.15. Heavy and light chains derived from the same cell were subsequently paired, and clonotypes were assigned based on their V and J genes using in-house R and Perl scripts. All scripts and the data used to process antibody sequences are publicly available on GitHub (https://github.com/stratust/igpipeline/tree/igpipeline2_timepoint_v2).

The frequency distributions of human V genes in anti-SARS-CoV-2 antibodies from this study was compared to 131,284,220 IgH and IgL sequences generated by (*50*) and downloaded from cAb-Rep (*51*), a database of human shared BCR clonotypes available at https://cab-rep.c2b2.columbia.edu/. Based on the 150 distinct V genes that make up the 1099 analyzed sequences from Ig repertoire of the 6 participants present in this study, we selected the IgH and IgL sequences from the database that are partially coded by the same V genes and counted them according to the constant region. The frequencies shown in fig. S3 are relative to the source and isotype analyzed. We used the two-sided binomial test to check whether the number of sequences belonging to a specific *IGHV* or *IGLV* gene in the repertoire is different according to the frequency of the same IgV gene in the database. Adjusted p-values were calculated using the false discovery rate (FDR) correction. Significant differences are denoted with stars.

Nucleotide somatic hypermutation and Complementarity-Determining Region 3 (CDR3) length were determined using in-house R and Perl scripts. For somatic hypermutations (SHM), *IGHV* and *IGLV* nucleotide sequences were aligned against their closest germlines using Igblastn and the number of differences were considered nucleotide mutations. The average number of mutations for V genes was calculated by dividing the sum of all nucleotide mutations across all participants by the number of sequences used for the analysis.

### Data presentation

Figures arranged in Adobe Illustrator 2022.

## Supplementary Figure Legends

**Fig. S1:**
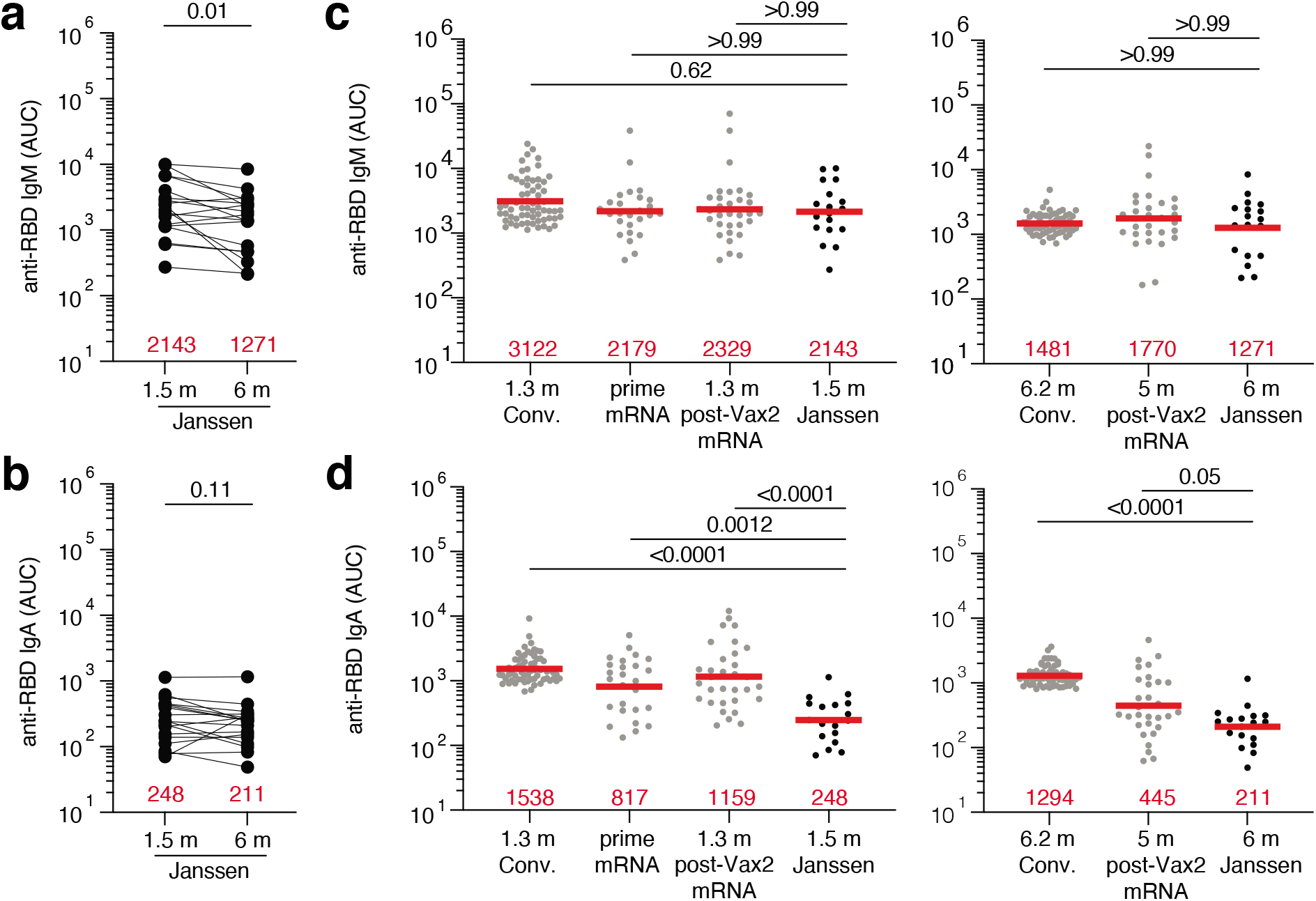
Plasma ELISA. **a-b,** Graph shows area under the curve (AUC) for **a,** plasma IgM and **b,** plasma IgA antibody binding to SARS-CoV-2 Wuhan-Hu-1 RBD after 1.5 months (m) and 6 m post-vaccination for n=18 samples. Lines connect longitudinal samples. **c-d,** Graph shows AUC for **c,** plasma IgM and **d,** plasma IgA binding to RBD in convalescent infected individuals 1.3 m post infection (*11*), and mRNA vaccinees after prime or 1.3 m post-second vaccination (Vax2) (*9*) compared to Janssen vaccinees 1.5 m post vaccination (left panel), or convalescent infected individuals 6.2 months post infection (*12*) and mRNA vaccinees 5 m post-Vax2 (*9*) compared to Janssen vaccinees at 6 m post vaccination (right panel). Red bars and values represent geometric mean values. Statistical significance in **a,** and **b,** was determined by Wilcoxon matched-pairs signed rank test. **c,** and **d,** was determined by two-tailed Kruskal-Wallis test with subsequent Dunn’s multiple comparisons.

**Fig. S2:**
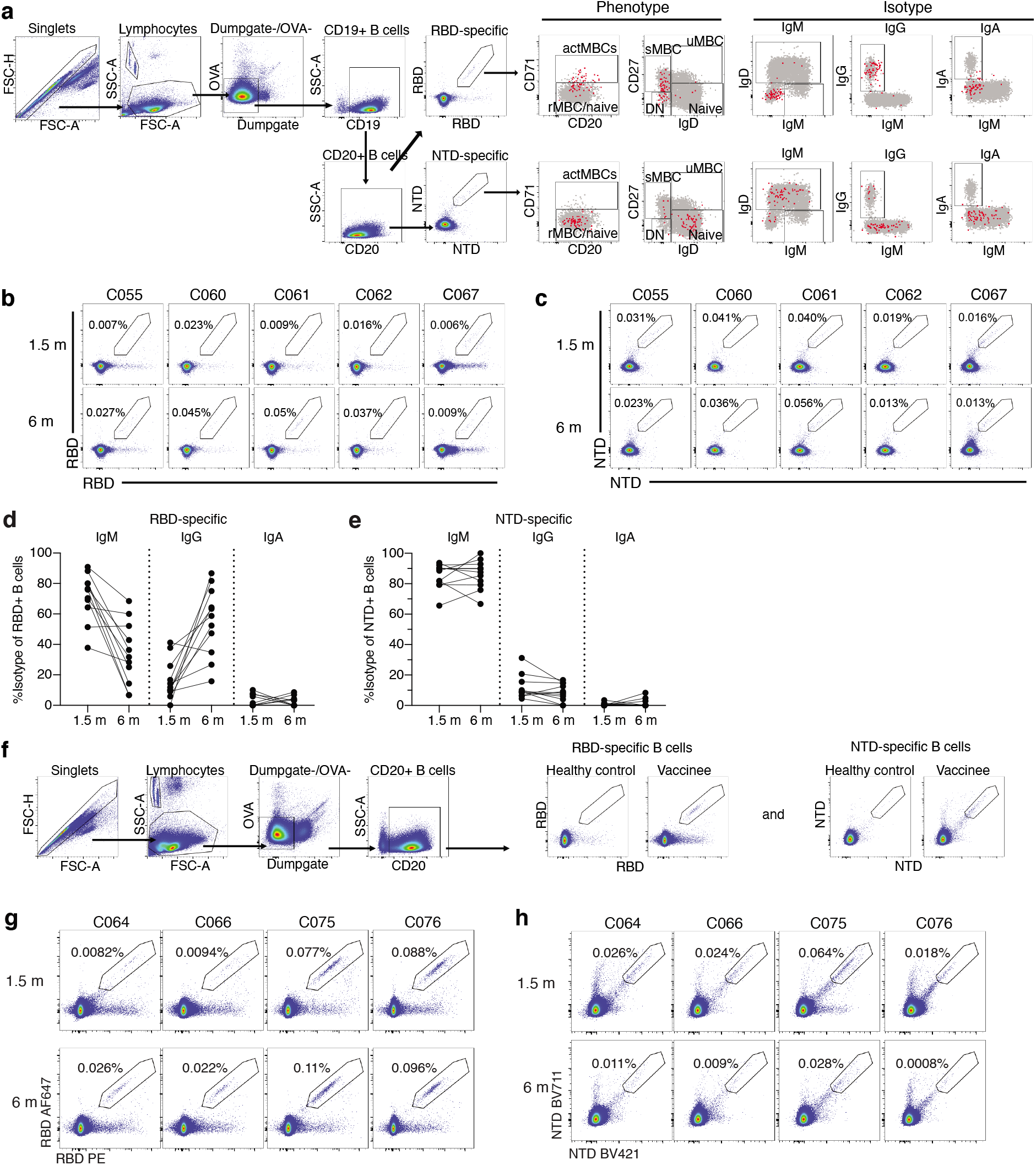
Flow Cytometry. **a,** Gating strategy for phenotyping. Gating was on lymphocytes singlets that were CD19^+^ or CD20^+^ and CD3-CD8-CD16-Ova-. Anti-IgG, IgM, IgA, IgD, CD71 and CD27 antibodies were used for B cell phenotype analysis. Antigen-specific cells were detected based on binding to Wuhan-Hu-1 RBD-PE^+^ and RBD-AF647^+^, or to Wuhan-Hu-1 NTD-BV711+ and NTD-BV421+. Counting beads were added to each sample and gated based on forward scatter (FSC) and side scatter (SSC) as per manufacturer instructions. **b-c,** Representative flow cytometry plots of **b,** RBD-binding B cells or **c,** NTD-binding B cells in 5 individuals after 1.5- and 6-months post vaccination. **d-e,** Graph showing the frequency of IgM, IgG, and IgA isotype in **d,** RBD-specific B cells and **e,** NTD-specific B cells after 1.5- or 6-months post-vaccination. **f,** Gating strategy for single-cell sorting for CD20+ B cells for Wuhan-Hu-1 RBD-PE and RBD-AF647 or Wuhan-Hu-1 NTD-BV711 and NTD-BV421. **g-h,** Representative flow cytometry plots showing **g,** dual AlexaFluor-647- and PE-Wuhan-Hu-1-RBD binding and **h,** BrilliantViolet-711- and BrilliantViolet-421-Wuhan-Hu-1 NTD binding, single-cell sorted B cells from 4 additional individuals at 1.5 months (m) or 6 m after vaccination. Percentage of antigen-specific B cells is indicated.

**Fig. S3:**
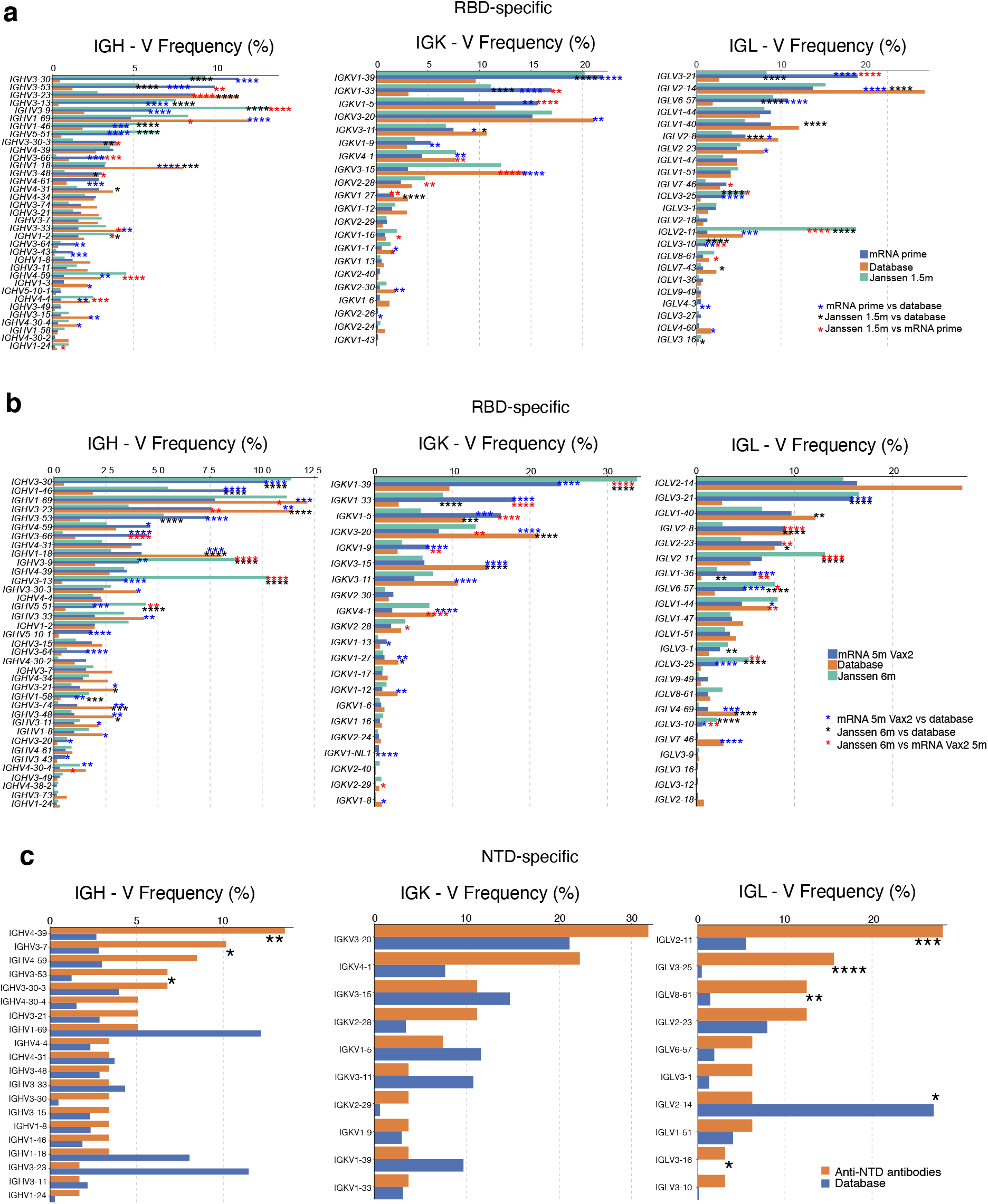
Frequency distribution of human V genes in SARS-CoV-2 RBD- and NTD-binding B cells. **a-b,** Comparison of the frequency distribution of human V genes for heavy chain and light chains of anti-RBD antibodies from this study and from a database of shared clonotypes of human B cell receptor generated by Cinque Soto et al (*50*). Graph shows relative abundance of human *IGHV* (left panel), *IGKV* (middle panel) and *IGLV* (right panel) genes in Sequence Read Archive accession SRP010970 (orange), Janssen antibodies (green), and mRNA vaccinees (blue), comparing **a,** 1.5 months (m) after Janssen vaccination to 1.3 months after one dose of mRNA vaccine (prime), or **b,** 6m post-Janssen vaccination to 5m after second dose of mRNA vaccine. Statistical significance was determined by two-sided binomial test. * = p≤0.05, ** = p≤0.01, *** = p≤0.001, **** = p≤0.0001. Color of stars indicates: black – comparing Janssen vaccination vs human database; blue – comparing mRNA vaccination vs human database; red - Janssen vaccination vs mRNA vaccination. **c,** Comparison of the frequency distribution of human V genes for heavy chain and light chains of all anti-NTD antibodies from this study to a database of shared clonotypes of human B cell receptor generated by Cinque Soto et al (*50*). Graph shows relative abundance of human *IGHV* (left panel), *IGKV* (middle panel) and *IGLV* (right panel) genes in Sequence Read Archive accession SRP010970 (blue), Janssen antibodies (orange). Statistical significance was determined by two-sided binomial test. * = p≤0.05, ** = p≤0.01, *** = p≤0.001, **** = p≤0.0001.

**Fig. S4:**
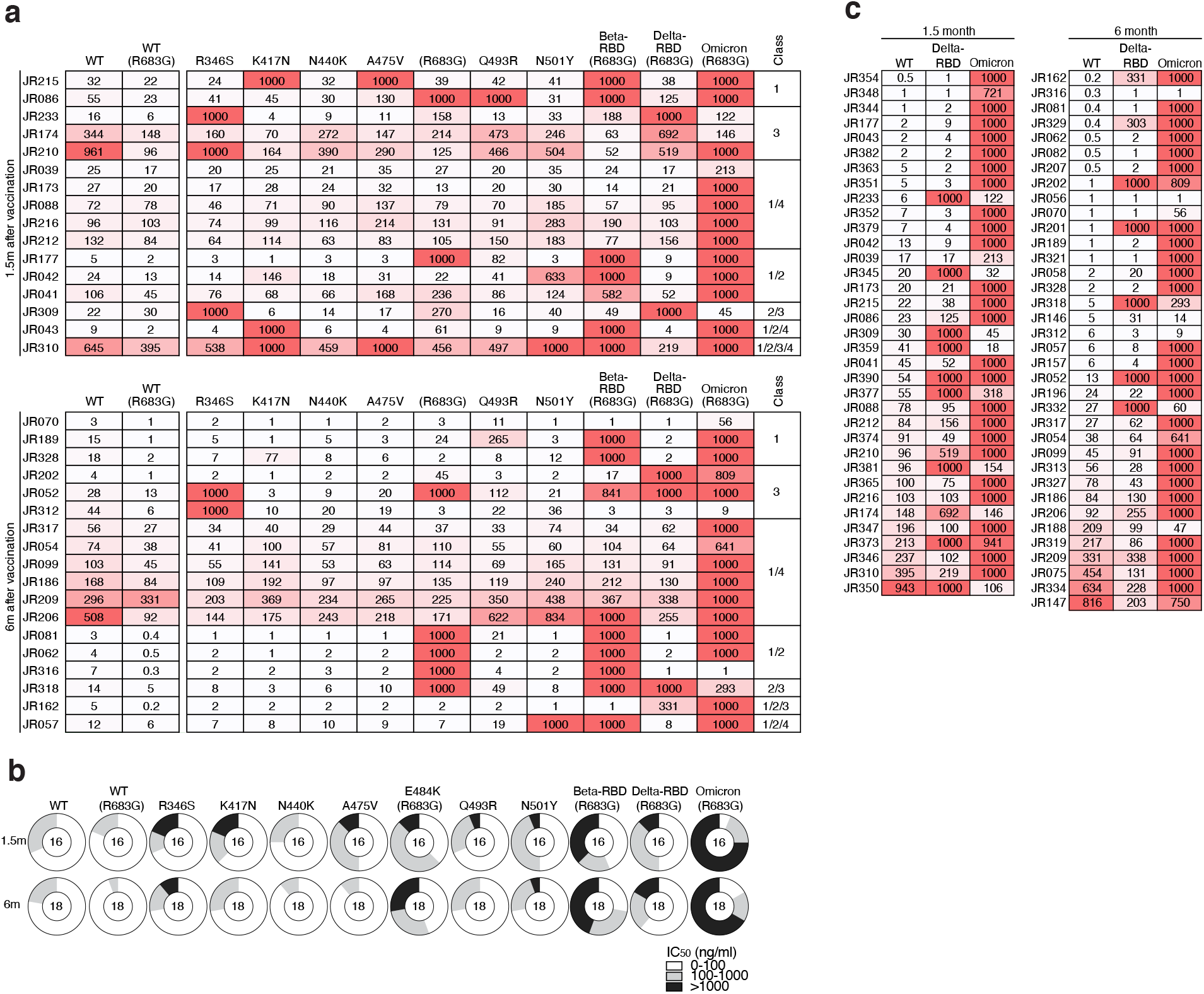
Neutralizing Breadth. **a,** Heat-maps show IC_50_s of antibodies shown in Fig. 5a against indicated mutant SARS-CoV-2 pseudoviruses listed across the top. Heatmap ranging from 0.1-1000 ng/ml in white to red. Antibody Classes are listed to the right, and were determined by competition BLI (see Fig. 4). **b,** Ring plots showing fraction of mAbs shown in Fig. 5a determined to be potently neutralizing (IC_50_ 1-100 ng/mL, white), poorly neutralizing (IC_50_ 100-1000 ng/mL, grey), or non-neutralizing (IC_50_>1000 ng/mL, black). Mutant or variant SARS-CoV-2 pseudovirus tested is indicated across the top and time point to the left. The number inside the circle indicated the number of antibodies tested. **c,** Heat map of antibodies shown in Fig. 5b, showing IC_50_s of antibodies detected at 1.5 months post vaccination (left panel, n=35) or 6 months post vaccination (right panel, n=36), against indicated variant SARS-CoV-2 pseudovirus listed across the top. Heatmap ranging from 0.1-1000 ng/ml in white to red. The E484K, K417N/E484K/N501Y and L452R/T478K substitution, as well as the deletions/substitutions corresponding to viral variants were incorporated into a spike protein that also includes the R683G substitution, which disrupts the furin cleavage site and increases particle infectivity. Neutralizing activity against mutant pseudoviruses were compared to a wildtype (WT) SARS-CoV-2 spike sequence (NC_045512), carrying R683G where appropriate.

